## Supplementary File 1 for "A processive rotary mechanism couples substrate unfolding and proteolysis in the ClpXP degradation machinery"

**Table 1.** Cryo-EM data acquisition, processing, atomic model statistics, and map/model depositions.

A. Cryo-EM data acquisition and image processing.

| **Data Collection** | | |
|---|---|---|
| | NmClpXP + GFP-SsrA | apo-NmClpP |
| Electron Microscope | Titan Krios | Tecnai F20 |
| Camera | Falcon 3EC | K2 Summit |
| Voltage (kV) | 300 | 200 |
| Nominal Magnification | 75,000 | 25,000 |
| Calibrated physical pixel size (Å) | 1.06 | 1.45 |
| Total exposure (e/Å$^2$) | 42.7 | 35 |
| Exposure rate (e/pixel/s) | 0.8 | 5 |
| Number of frames | 30 | 30 |
| Defocus range (μm) | 0.9 to 1.7 | 0.8 to 3.3 |
| **Image Processing** | | |
| Motion correction software | *cryoSPARC v2* | *cryoSPARC v2* |
| CTF estimation software | *cryoSPARC v2* | *cryoSPARC v2* |
| Particle selection software | *cryoSPARC v2* | *cryoSPARC v2* |
| Micrographs used | 2,680 | 122 |
| Particle images selected | 466,549 | 100,132 |
| 3D map classification and refinement software | *cryoSPARC v2* | *cryoSPARC v2* |

B. Map and model statistics.

| **EM maps** | NmClpXP with Symmetry | Focused NmClpX Conformation A | Focused NmClpX Conformation B | Apo-NmClpP |
|---|---|---|---|---|
| Particle images contributing to maps | 377,234 | 110,696 | 178,448 | 50,403 |
| Applied symmetry | D7 | C1 | C1 | D7 |
| Applied B-factor (Å$^2$) | -116.2 | -84.1 | -88.1 | -229.7 |
| Global resolution (FSC = 0.143, Å) | 2.3 | 3.3 | 2.9 | 4.1 |

| **Model Building** | NmClpX Conformation A focused | NmClpX Conformation B focused | NmClpP with Symmetry | NmClpXP Conformation A combined | NmClpXP Conformation B combined |
|---|---|---|---|---|---|
| Modeling software | Coot, Phenix, Rosetta | | | | |
| Number of residues | 1912 | 2018 | 2671 | 4583 | 4689 |
| RMS bond length (Å) | 0.0043 | 0.0037 | 0.0039 | 0.0044 | 0.0038 |
| RMS bond angle (º) | 1.04 | 0.93 | 0.91 | 1.00 | 0.92 |
| Ramachandaran outliers (%) | 0.00 | 0.05 | 0.00 | 0.00 | 0.02 |
| Ramachandran favoured (%) | 98.33 | 97.32 | 95.68 | 97.07 | 96.39 |
| Rotamer outliers | 0.25 | 0.24 | 0.00 | 0.13 | 0.10 |
| C-beta deviations | 0 | 0 | 0 | 0 | 0 |
| Clashscore | 1.45 | 2.8 | 0.67 | 1.84 | 2.11 |
| MolProbity score | 0.88 | 1.20 | 1.02 | 1.11 | 1.22 |
| EMRinger score | 2.78 | 3.42 | 5.6 | | |
| Map-Model CC_mask | 0.75 | 0.78 | 0.82 | | |
| Ligand | 2 ADP<br>4 Mg-ATP | 1 ADP<br>5 Mg-ATP | | 2 ADP<br>4 Mg-ATP | 1 ADP<br>5 Mg-ATP |

C. Residues excluded in atomic models*.

| **ClpX Protomer** | NmClpXP Conformation A combined | NmClpXP Conformation B combined |
|---|---|---|
| X1 (Chain A) | 1-62, 102-109, 149-156, 191-204, 224-236, 263-264, 273-282, 413-414 | 1-62, 228-233, 275-278, 413-414 |
| X2 (Chain B) | 1-62, 102-111, 192-198, 226-235, 275-279, 413-414 | 1-62, 226-233, 275-279, 413-414 |
| X3 (Chain C) | 1-62, 192-198, 225-232, 275-279, 413-414 | 1-62, 192-198, 228-233, 274-278, 413-414 |
| X4 (Chain D) | 1-62, 103-106, 193-198, 226-234, 276-280, 413-414 | 1-62, 193-198, 229-233, 413-414 |
| X5 (Chain E) | 1-62, 193-198, 229-233, 277-281, 413-414 | 1-62, 193-198, 277-279, 413-414 |
| X6 (Chain F) | 1-62, 193-201, 225-234, 259-287, 413-414 | 1-62, 191-204, 227-233, 275-282, 413-414 |

* Protomer X1 is US in Conformation A and protomer X2 is LS in Conformation B

| **ClpP subunits** | NmClpXP Conformation A combined | NmClpXP Conformation B combined |
|---|---|---|
| Chain H | 1-5, 16-18, 199-204 | 1-5, 16-18, 199-204 |
| Chain I | 1-5, 16-17, 199-204 | 1-5, 16-17, 199-204 |
| Chain J | 1-5, 17, 199-204 | 1-5, 17, 199-204 |
| Chain K | 1-5, 199-204 | 1-5, 199-204 |
| Chain L | 1-5, 16-17, 199-204 | 1-5, 16-17, 199-204 |
| Chain M | 1-5, 13-19, 199-204 | 1-5, 13-19, 199-204 |
| Chain N | 1-5, 16-17, 199-204 | 1-5, 16-17, 199-204 |
| Chain O | 1-5, 16-17, 199-204 | 1-5, 16-17, 199-204 |
| Chain P | 1-5, 16-17, 199-204 | 1-5, 16-17, 199-204 |
| Chain Q | 1-5, 16-17, 199-204 | 1-5, 16-17, 199-204 |
| Chain R | 1-5, 16-17, 199-204 | 1-5, 16-17, 199-204 |
| Chain S | 1-5, 16-17, 199-204 | 1-5, 16-17, 199-204 |
| Chain T | 1-5, 16-17, 199-204 | 1-5, 16-17, 199-204 |
| Chain U | 1-5, 16-17, 199-204 | 1-5, 16-17, 199-204 |

D. Deposited maps and associated coordinate files.

| **Maps** | **EMDB code** | **Associated PDB ID** |
|---|---|---|
| NmClpXP Conformation A | EMD-XXXX | XXXX |
| NmClpXP Conformation B | EMD-XXXX | XXXX |
| NmClpXP D7 | EMD-XXXX | |
| Apo-NmClpP | EMD-XXXX | |
